# Mavacamten improves energy balance in a pre-clinical model of RASopathy-associated hypertrophic cardiomyopathy

**DOI:** 10.1101/2025.02.21.639222

**Authors:** Andrea Ruiz-Velasco, Charlène Jouve, Lucille Deshayes, Michael Kohlhaas, Christoph Maack, Jean-Sébastien Hulot

**Author notes:** **Address for correspondence:** Prof. Jean-Sébastien Hulot, PARCC, 56 Rue Leblanc, F-75015 Paris, France.

## Abstract

**Background and aim:** Developmental disorders caused by activating mutations in the RAS-MAPK pathway account for nearly 20% of hypertrophic cardiomyopathy (HCM) cases in paediatric patients. Compared to sarcomeric HCM, RAS-HCM presents a higher risk of obstruction and hospitalisation. The myosin inhibitor mavacamten has been approved in the European Union for treating adults with obstructive HCM; however, clinical trials have excluded syndromic HCM. Consequently, this study aimed to characterise the functional and energetic disturbances induced by the RASopathy mutation BRAF p.Thr599Arg in cardiomyocytes and to evaluate the effects of mavacamten treatment.

**Methods:** Human induced pluripotent stem cell-derived cardiomyocytes (hiPSC-CMs) with a CRISPR-induced BRAF T599R mutation and their isogenic control were employed to model RAS-HCM in vitro. The cell size, contractility, and transcriptomic profile were assessed to determine the phenotype of the cardiomyocytes. Energetics were evaluated using the Mito Stress assay, live ATP levels, and NAD(P)H and FAD+ autofluorescence.

**Results:** BRAF-mutant cardiomyocytes demonstrated hypertrophy and hypercontractility. Furthermore, energetic profiling revealed increased mitochondrial NAD(P)H and FAD+ pools and an enhanced energetic state in the Mito Stress assay with increased maximal respiratory capacity. However, they also exhibited a significant ATP drop during rapid pacing compared to the control, suggesting that mitochondrial capacity remains insufficient to meet the ATP demand. Mavacamten treatment normalised excessive ATP consumption during acute pacing, suggesting reduced mitochondrial overactivity.

**Conclusions:** BRAF-mutant cardiomyocytes recapitulate the characteristics of HCM in vitro. Mavacamten mitigates dysfunctions and restores energetic balance under stress conditions, indicating it holds potential as a therapeutic option for RASopathy-associated HCM.

## Introduction

Hypertrophic cardiomyopathy (HCM) is a common genetic heart disease characterised by left ventricular hypertrophy, hypercontractility, and susceptibility to arrhythmias, heart failure, and sudden cardiac death. It is estimated to have a global prevalence of 1 in 500 individuals (1) and is generally attributed to mutations in sarcomeric genes, typically *MYH7* and *MYBPC3 (2)*. While sarcomeric protein mutations dominate the cause in adult cases, up to 18% of pediatric HCM cases stem from RASopathies (3, 4), a group of developmental disorders caused by germline mutations in the RAS-MAPK signalling pathway. RASopathies, including Noonan syndrome and cardiofaciocutaneous syndrome (CFCS), and the prevalence of HCM in these groups has been reported to be as high as 35-40% (5-9). RASopathies are associated with a spectrum of cardiac and systemic abnormalities, posing significant and unique therapeutic challenges.

RASopathy-associated HCM (RAS-HCM) mimics the HCM phenotype and even shares many of the underlying mechanisms, explaining why RAS-HCM is often considered a phenocopy of HCM. However, its progression and, therefore, its management are currently different. Hypercontractility is a predominant pathological mechanism in HCM, driving cardiomyocyte growth, metabolic remodelling, oxidative stress, and susceptibility to arrhythmias (10). More specifically, in RAS-HCM, the activation of the RAS-MAPK pathway induces hypertrophy through the ERK1/2 signalling cascade and can affect diastolic function by modulating calcium transients and uptake (11, 12). Additionally, changes in contraction force and sarcomeric disarray have also been observed (13, 14), but the molecular mechanism linking RAS-MAPK signalling to changes in the contractile apparatus has not been fully described.

In RAS-HCM, there is a high prevalence and severity of left ventricular outflow tract obstruction and higher rate of heart failure hospitalizations and mortality compared to HCM from other etiologies (4, 5, 15, 16). Current recommended treatment includes the use of beta-blockers and selective management of obstructions (17). The use of MEK inhibitors has shown some promise for the reversal of cardiac hypertrophy but requires further development (18). Mavacamten, a selective inhibitor of cardiac myosin ATPase, has emerged as a targeted therapy for HCM and was recently approved in the European Union for the treatment of adults with obstructive HCM (19). By targeting sarcomeric hypercontractility, it improves myocardial relaxation and reduces left ventricular outflow tract obstruction (20). Its use in the context of RASopathies is an area of emerging interest due to the potential overlap in the pathophysiology with other forms of HCM. However, clinical trials on mavacamten, including pivotal trials like EXPLORER-HCM (21), have excluded patients with secondary or syndromic HCM, and the therapeutic potential of mavacamten in non-sarcomeric forms of HCM, including RAS-HCM, has therefore not been demonstrated.

This study sought to address this gap by examining an in vitro model of RASopathy-associated HCM using human induced pluripotent stem cells (hiPSC) – derived cardiomyocytes with a BRAF p.Thr599Arg activating mutation (BRAF T599R). We characterized the functional, morphological, and energetic abnormalities in these mutant cardiomyocytes and evaluated the effects of mavacamten treatment on these pathological features.

## Methods

### hiPSC culture and genome editing with CRISP/Cas9

The two hiPSC cell lines employed here have been previously reported. Briefly, the control cell line SKiPSC-31-3 was obtained from a healthy male adult donor (22-25). The *BRAF* p.Thr663Arg mutation was introduced by CRISPR-Cas9 homology-directed repair (HDR) with the guide RNA and single-stranded oligodeoxynucleotides (ssODN) template sequences listed below (26). Briefly, the sgRNA-Cas9 plasmid (Addgene plasmid #62988) and ssODN were introduced into hiPSCs by electroporation and transfected cells were enriched by puromycin selection. Single-round colonies were amplified for Sanger sequencing (Eurofins genomic), and a homozygote clone was selected for this project. Cell pluripotency was assessed by quantitative PCR and immunofluorescence (Supplementary Figure 1), and genome stability was evaluated by detecting recurrent genetic abnormalities (iCS-digital PSC test, Stem Genomics) (27). HiPSC were kept on Matrigel-coated (Corning) dishes with mTeSR Plus medium (STEMCell Technologies) at 37 °C with 5% CO2 atmosphere. They were passaged manually once a week by scrapping with a pipette tip. Cells were routinely screened for mycoplasma with the MycoAlert kit (Lonza).

- *sgRNA BRAF:* TCGAGATTTCACTGTAGCTA
- *ssODN T599R:* GGCCAAAAATTTAATCAGTGGAAAAATAGCCTCAATTCTTACCAT CCACAAAATGGATCCAGACAACTGTTCAAACTGATGGGACCCACTCCATCGAGATTTCACT CTCGCTAGACCAAAATCACCTAT
- *Fwd-BRAF:* CCTAACACATTTCAAGCCCCA
- *Rev-BRAF:* AACTCAGCAGCATCTCAGGG

### Cardiomyocyte differentiation

For cardiomyocyte differentiation, hiPSC were dissociated into small aggregates with ReLeSR (STEMCell Technologies) and seeded on Matrigel-coated plates. At approximately 90% confluency, the mTeSR™ Plus medium was replaced with RPMI-1640 (Thermo Fisher Scientific) supplemented with B27 minus insulin (Thermo Fisher Scientific) and 6 µM CHIR-99021 (Abcam). Cells were maintained under a 5% CO_2_ atmosphere for 48 hours. The medium was then changed to RPMI-1640/B27 minus insulin (Thermo Fisher Scientific) for 24 hours, followed by RPMI-1640/B27 minus insulin supplemented with 5 µM IWR-1 (Sigma-Aldrich) for 48 hours. From day 5 to day 7, cells were cultured in RPMI-1640/B27 minus insulin. On day 7, the medium was switched to RPMI-1640 with B27 plus insulin (Thermo Fisher Scientific), and regular media changes continued. Beating cardiomyocytes were observed around day 9 and by day 11 they were subjected to a three-day glucose starvation in RPMI-B27 without glucose (Thermo Fisher Scientific). After cells were dissociated using 0.05% trypsin (Thermo Fisher Scientific) for 10 minutes and seeded at 0.3 × 106 cells/cm2 in RPMI-1640/B27 plus insulin for 24 hours before a second round of glucose starvation. On day 18, cells were cultured in RPMI-1640/B27 plus insulin with medium changes every two days. All analyses were performed on cells 35 days post-differentiation.

### Basal cell size and mitochondrial ATP measurement

Cell cross-sectional area and basal ATP were measured on live cardiomyocytes seeded on Matrigel-coated glass-bottom 3.5 mm dishes. The ATP tracker (BioTracker ATP-Red Live Cell Dye, Millipore) 10 mM stock solution was diluted to a 5 µM concentration in RPMI-B27 medium and warmed to 37 °C. The cell medium was replaced with the ATP tracker medium, and the cells were placed back in the incubator for 20 minutes. In the meantime, 5 µM wheat germ agglutinin-fluorescein (Vector Laboratories) was prepared in Hank’s balanced salt solution (Thermo Fisher Scientific) and heated to 37 °C. The ATP tracker medium was replaced with the WGA solution and incubated for 10 mins at 37 °C. At the end of the incubation, the medium was changed back to warm RPMI-B27 medium. Fluorescent images were acquired using a Leica TSC SP8 confocal microscope (Leica Microsystems) with a heated chamber and processed with Fiji (28). Individual cell areas were drawn manually on the WGA channel, and the same regions were used to measure the ATP mean fluorescence intensity for each cardiomyocyte.

### Live ATP monitoring with electrical pacing

For the electrical stimulation, cardiomyocytes were kept on Matrigel-coated 12-well plates. On the day of the imaging, they were labelled by incubating with a 5 µM ATP tracker prepared in RPMI-B27 medium for 20 mins at 37 °C. Then, this was replaced by a warm RPMI-B27 medium supplemented with 1.8 mM CaCl_2_ (Thermo Fisher Scientific) and 0.1 mM pyruvate (Thermo Fisher Scientific). The C-dish electrode plate (Ionoptix) was placed on top of the plate, covered with transparent adhesive film, and connected to the C-pace EM module (Ionoptix). The sequential stimulation used 4 ms pulses at 5 V: 1 Hz for 5 minutes, 1.5 Hz for 5 minutes, 2 Hz for 20 minutes. Fluorescent images were acquired using a Leica TSC SP8 confocal microscope (Leica Microsystems) with a heated chamber at 4-minute intervals. The whole image’s raw intensity of fluorescence was measured with Fiji (28). Fractional differences and areas under the curves were calculated with GraphPad Prism 10.4.1 Software.

### Flow cytometry

Single-cell suspensions of hiPSC-derived cardiomyocytes were obtained using 0.05% trypsin-EDTA (Thermo Fisher Scientific) and stained with Zombie NIR™ viability dye (BioLegend) following the manufacturer’s instructions. Cells were fixed and permeabilised with the Inside Stain Kit (Miltenyi Biotec) for 10 mins at room temperature, followed by a 10-minute incubation with cTnT-APC or control IgG1-APC REAfinity antibody (Miltenyi Biotech). Data was acquired with a BD LSR Fortessa X-20 (BD Biosciences) flow cytometer. At least 30,000 cells were recorded, and single-labelled controls were used to calculate the compensation. Results were analysed using FlowJo v10 software (FlowJo, LLC).

### RNA-sequencing and gene set enrichment analysis (GSEA)

RNA was extracted with the NucleoSpin RNA Kit (Macherey-Nagel) according to the manufacturer’s instructions and quantified using a NanoDrop (Thermo Fisher Scientific). Integragen prepared the RNA-sequencing libraries with the NEBNext® UltraTM II mRNA-Seq kit. Paired-end sequencing was performed with the Illumina NovaSeq6000 S2. STAR alignment and FPKM quantification were performed using Gencode v31 annotation (restricted to protein-coding genes, antisense and lincRNAs). Raw counts for each sample were imported into R statistical software. The extracted count matrix was normalised for library size and coding length of genes to compute FPKM expression levels.

#### Unsupervised analysis

The Bioconductor edgeR package was used to import raw counts into R statistical software and compute normalised log2 CPM (counts per millions of mapped reads) using the TMM (weighted trimmed mean of M-values) as a normalisation procedure. The normalised expression matrix from the 1000 most variant genes (based on standard deviation) was used to classify the samples according to their gene expression patterns using principal component analysis (PCA), hierarchical clustering and consensus clustering. tSNE analysis was performed with the Bioconductor Rtsne package.

#### Differential expression analysis

Differential expression analysis was performed using the Bioconductor limma package and the voom transformation. To improve the statistical power of the analysis, only genes expressed in at least one sample were considered. A qval threshold of <= 0.05 and a minimum fold change of 1.2 were used to define differentially expressed genes.

#### Pathway enrichment analysis – GSEA

The gene list from the differential analysis was ordered by decreasing log2 fold change. Gene set enrichment analysis was performed by clusterProfiler::GSEA function using the fgsea algorithm. Gene sets from the MSigDB v7.4 database were selected among the C2_curated, C5_GO, and Hallmark classes, keeping only gene sets defined by 10-500 genes.

### Contractility assessment

Monolayer contractility was evaluated with the MuscleMotion (29) plugin for Fiji (28). Twenty-second uncompressed AVI videos were recorded with a high-speed CCD camera (PL-D672MU, Pixelink) in a Zeiss Primovert microscope. Three regions were recorded per well and averaged.

### Redox index and mitochondrial NAD(P)H and FAD+ pool estimation

Cells were perfused with a warm Tyrode solution containing 130 mM NaCl, 5 mM KCl, 1 mM MgCl_2_, 2 mM CaCl_2_, 10 mM Na-HEPES, 10 mM glucose, 2 mM Na-pyruvate, and 0.3 mM ascorbic acid, pH 7.4. They were paced by electrical field stimulation using a customised IonOptix system, as described previously (30). The autofluorescence of NAD(P)H/NAD(P)^+^ and FADH_2_/FAD were determined by alternately exciting cells at 340 and 485 nm, and collecting emission at 450 and 525 nm, respectively. Calibration was performed with FCCP (5 μM) and cyanide (4 mM). The redox index was estimated from the calibrated values. The mitochondrial pools of mutant cardiomyocytes are expressed relative to the control.

### Mitochondrial respiration and analysis

hiPSC-CMs were cultured on custom micropatterned coverslips (18 mm rounds, 30 µm-wide adhesive lines alternated with non-adhesive regions; 4D cell). Coverslips were coated with Matrigel (Corning), and cells were seeded on day 35 at a 12.5 × 10^4^ cells/cm^2^ density. Cultures were maintained in RPMI-1640+B27 plus (Thermo Fisher Scientific) for 7 days at 37°C in a 5% CO_2_ environment.

For the bioenergetic analysis, hiPSC-cardiomyocytes were seeded in Matrigel-coated Seahorse XF96 plates (Agilent Technologies) and subjected to extracellular flux analysis. According to manufacturer instructions, mitochondrial respiration (OCR) was measured using oligomycin, FCCP, and rotenone/antimycin A (Sigma-Aldrich). Each differentiation was measured in quadruplicates. Data were normalised to protein content determined via Bradford assay (Bio-Rad).

### Immunofluorescence

Cardiomyocytes were fixed with 4% paraformaldehyde (Electron Microscopy Sciences) for 10 minutes, permeabilised with 0.5% Triton X-100 (Sigma-Aldrich), and blocked in a solution of 2% bovine serum albumin (Sigma-Aldrich) for one hour. Cells were incubated overnight with the primary antibodies (Table below) at 4°C, followed by Alexa Fluor-conjugated secondary antibodies (Thermo Fisher Scientific) for one hour at room temperature. Nuclei were counterstained with DAPI (Sigma-Aldrich), and coverslips were mounted with Dako Faramount Mounting Medium (Agilent Technologies). Fluorescent images were acquired using a Leica TSC SP8 confocal microscope (Leica Microsystems) and processed with Fiji software.

| Antibody | Catalogue no. | Manufacturer | Dilution |
| --- | --- | --- | --- |
| cTnT | ab45932/MA. 5-12960 (Rb) | Abcam/ThermoFisher | 1:1000 |
| α-Actinin | A7811 | Sigma-Aldrich | 1:1000 |
| MLC2V | 10906-1-AP | Proteintech | 1:600 |
| MLC2A | 311011 | Synaptic Systems | 1:1000 |
| Tra1-60 | MAB4360 | Millipore | 1:100 |
| Nanog | 4903 | Cell Signaling | 1:200 |
| Tra1-81 | MAB4381 | Millipore | 1:100 |
| Sox2 | AB5603 | Millipore | 1:200 |
| Goat anti-Rabbit IgG (H+L), Alexa Fluor 546 | A11010 | Thermo Fisher Scientific | 1:1000 |
| Goat anti-Mouse IgG (H+L), Alexa Fluor 488 | A10680 | Thermo Fisher Scientific | 1:1000 |
| Goat anti-Rabbit IgG (H+L), Alexa Fluor 488 | A11008 | Thermo Fisher Scientific | 1:1000 |
| Goat anti-Mouse IgG (H+L), Alexa Fluor 546 | A11030 | Thermo Fisher Scientific | 1:1000 |

## Results

### BRAF T599R cardiomyocytes are hypertrophic and hypercontractile

Isogenic control iPSC from a healthy male donor and homozygote BRAF-mutant iPSC were previously described (26), but their pluripotency markers were validated here again (Supplementary Figure 1A). Cardiomyocytes were derived via the temporal modulation of the Wnt signaling pathway and enriched by glucose starvation (31, 32). Cardiac markers and spontaneous contraction were observed, and the differentiation efficiency of each batch was determined by flow cytometry (Supplementary Figure 1B-C). To evaluate cell size, the cross-sectional area of cardiomyocytes was measured by labelling the membrane and live cell imaging after 35 days in culture, and BRAF-mutant cardiomyocytes showed significant cell hypertrophy as compared to the isogenic controls (4224 vs 2797 μm^2,^ respectively, p-value 0.0080, Figure 1A). This result was confirmed by using the forward scatter area (FSC-A) to determine the cell size of the BRAF-mutant CMs relative to the control (Figure 1B). Gene expression analysis from bulk RNA sequencing did not detect clustering or segregation of the mutant CMs, and only three genes were differentially expressed when compared to the isogenic control (Supplementary Figure 2A-B). However, gene set enrichment analysis showed that BRAF T599R mutant cardiomyocytes have an upregulation of protein translation pathways, changes in the regulation of transcription, extracellular matrix and ion handling, characteristics previously reported for cardiac hypertrophy (33, 34). In addition, a trend towards increased expression can be observed in a panel of cardiac hypertrophy genes (Figure 1C-D).

**Figure 1.**
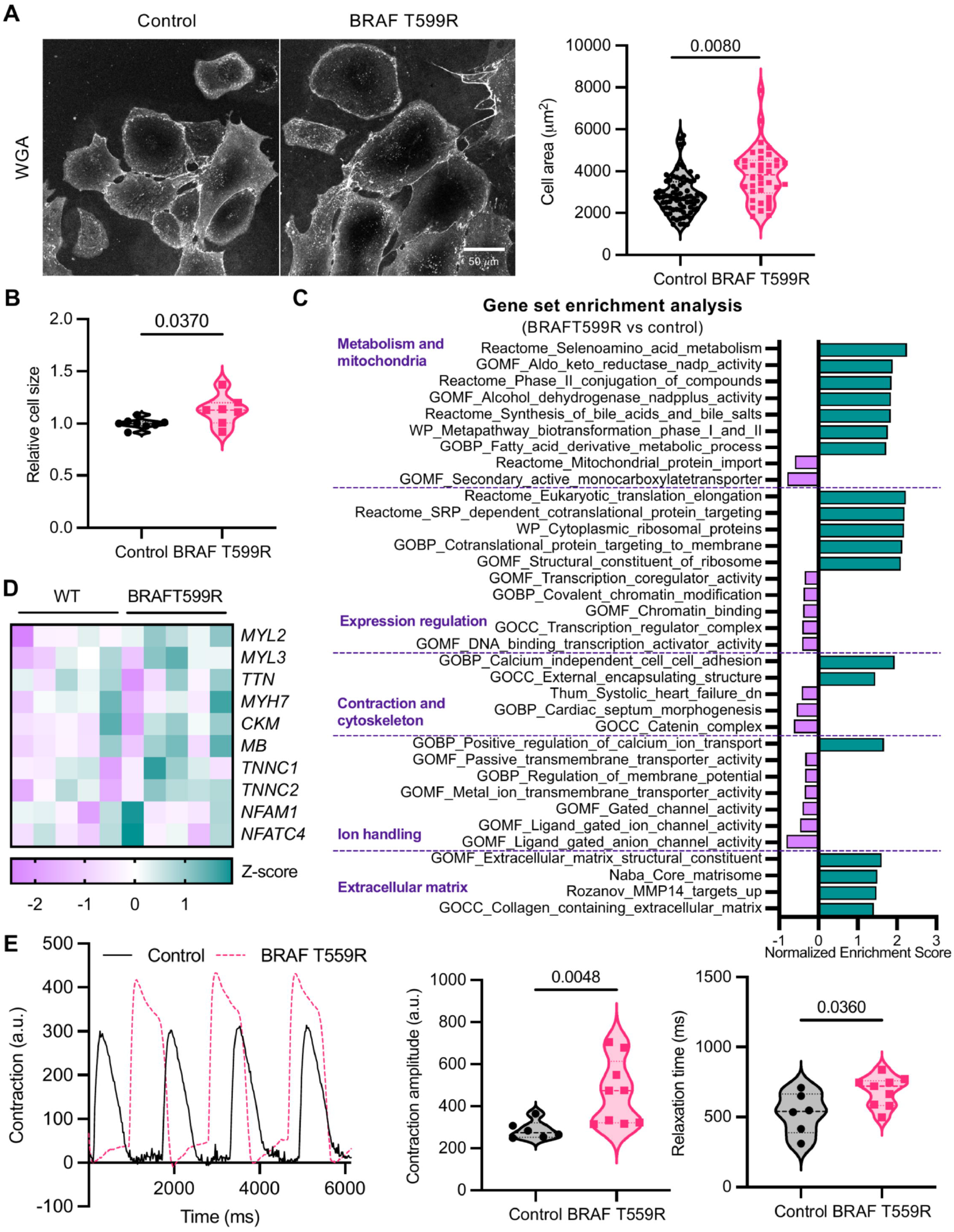
BRAFT599R cardiomyocytes are hypertrophic and hypercontractile. **A)** Representative confocal images of fluorescent wheat-germ aggluatinin (WGA) labeled control and BRAF-mutant cardiomyocytes (scale bar = 50 um) and the cell area quantification. N=8 differentiations (20-70 cells per differentiation). Nested-I test. B) Cell size determined by the mean forward scatter area (FSC-A) relative to the isogenic control using flow cytometry. N=7-8 differentiations (at least 5000 events per differentiation). C) Differentially expressed pathways in BRAF-mutant CMs compared to isogenic control identified by gene set enrichment analysis from bulk RNA-sequencing data performed with Galileo (lntegragen, France). The bar graph shows the normalized enrichment score colored in purple if underexpressed and in green if overexpressed (q-value<0.05). D). Heatmap showing genes involved in cardiac hypertrophy, in purple if underexpressed and in green if overexpressed. N=5 differentiations. E) Contraction profile of control and BRAF-mutant CM monolayers generated by MuscleMotion and violin plots of the contraction amplitude and relaxation time. N=6-9 differentiations. Violin plots show the distribution, median, and quartiles of the data. The statistical analysis used was a two-tailed Mann-Whitney test except where otherwise specified. p-value < 0.05 considered significant and displayed.

Hypercontractility is a feature of HCM; therefore, we recorded the spontaneous contractions of the cardiomyocytes and measured them using the MuscleMotion plugin for ImageJ (29). Control and BRAF-mutant cells showed similar contraction rates (Supplementary Table 1); however, the BRAF T599R activating mutation increased contraction amplitude and prolonged relaxation time (Figure 1E), showing a stronger contraction and impaired relaxation. Overall, these results indicate that the HCM phenotype associated with the activating BRAF T599R mutation can be recapitulated in vitro under baseline conditions.

### Acute stress reveals altered energetics in BRAF T599R cardiomyocytes

Hypertrophy and hypercontractility are high-energy demand processes. Next, we focused on characterising the energetic status of the cells. An ATP tracker was used to detect mitochondrial ATP in live cells. In basal conditions, BRAF-mutant cardiomyocytes showed no significant difference in ATP levels compared to the control (Figure 2A). Then, we applied electrical stimulation to pace the cardiomyocytes at increasing frequencies and monitor the ATP during the pacing and a short final recovery period (Figure 2B). We measured the area under the fluorescence change curve for ATP (AUC-ATP) to reflect the overall variation in ATP content. We observed a larger negative AUC-ATP in BRAF-mutant cardiomyocytes, indicating a substantial ATP drop in response to pacing than in the control cardiomyocytes. The difference was not significant for the recovery period, even if an increasing negative area can be observed (Figure 2C). Both cell lines contracted at the expected rate with the pacing (Supplementary Figure 3).

**Figure 2.**
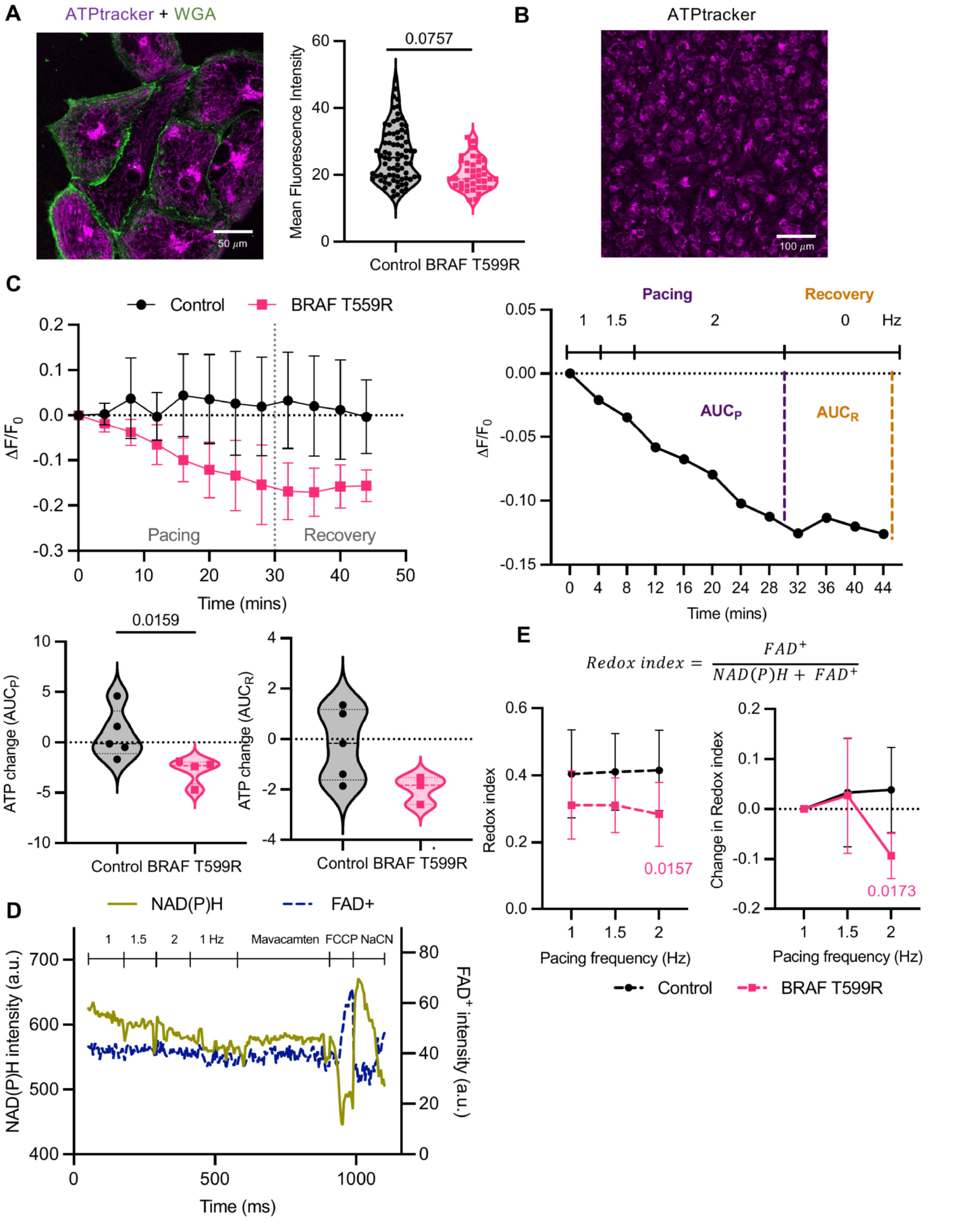
Acute stress reveals altered energetics in BRAFT599R cardiomyocytes. A) Confocal image of mitochondrial ATP labeled with fluorescent ATPtracker and the cell membrane with WGA marker (scale bar= 50 *µm)* and ATPtracker intensity quantification per cardiomyocyte. N=8 differentiations (20-70 cells per differentiation). Nested-I test. B) Wider field view used for live monitoring the ATPtracker using an incremental electrical pacing protocol and imaging every four minutes (scale bar = 100 *µm)*. The area under the normalized intensity curve (AUC) was measured for the pacing and recovery periods separately. C) Average ATPtracker fluorescence intensity changes from control and BRAF-mutant CMs and violin plots of the AUCs. N=4-5 differentiations. D) NAD(P)H and FAD+ autofluorescence measured during electrical pacing, mavacamtem (0.5 *µM)*, FCCP (5 *µM)*, and cyanide (4 mM) exposition. Data presented as mean ± standard deviation. E) The redox index was determined from the NAD(P)H and FAD+ intensities to represent the oxidative metabolism of the mitochondria. The fractional change in redox index relative to 1 Hz pacing is also shown. N=5 independent experiments from 2 differentiations. Two-way ANOVA with Dunnett’s multiple comparison test. Data presented as mean ± standard deviation. For others, violin plots show the data’s distribution, median, and quartiles. The statistical analysis used was a two-tailed Mann-Whitney test except where otherwise specified. p-value < 0.05 considered significant and displayed.

Additionally, we used NAD(P)H and FAD^+^ autofluorescence measurements to estimate mitochondrial metabolic activity during pacing as a redox index. These two cofactors are oxidised in the electron transport respiratory chain for ATP production; therefore, their ratio reflects the redox state of the mitochondria and changes in mitochondrial activity. For the measurements, the mitochondrial uncoupler, carbonylcyanide-p-trifluoromethoxyphenylhydrazone (FCCP), and the complex IV inhibitor, sodium cyanide (NaCN), were used to establish the minimum and maximum to calibrate each signal (Figure 2D). The redox ratio was defined as FAD^+^/[NAD(P)H+FAD^+^] (35), meaning that an increase in the ratio represents an increase in mitochondrial activity. In contrast, a decrease in the ratio represents a decreased mitochondrial activity. In basal conditions, control and BRAF-mutant cardiomyocytes show similar redox ratios; however, a significant decrease was observed in the BRAF-mutant cells in response to the 2 Hz pacing, suggesting a blunted mitochondrial activity (Figure 2E, Supplementary Figure 4). Overall, these results indicate that BRAF-mutant cardiomyocytes are not able to preserve their energetic balance in response to pacing.

### Energetic disturbances in BRAF T599R cardiomyocytes are not due to a decreased mitochondrial capacity

The energy deficiency detected with acute stress in BRAF-mutant cardiomyocytes could be due to a reduced mitochondrial mass or mitochondrial capacity. We estimated the maximally oxidisable mitochondrial pool for each cofactor from the NAD(P)H and FAD^+^ measurements. We observed that both pools are increased in the BRAF-mutant cardiomyocytes compared to the control, suggesting that the mitochondrial content or size is rather increased (Figure 3A-B). Thus, we proceeded to evaluate their mitochondrial respiration and capacity with the Seahorse MitoStress test. The oxygen consumption rate measurements (OCR) with complex inhibitors and mitochondrial uncoupler indicated that BRAF-mutant cardiomyocytes display a more energetic phenotype compared to the control. The BRAF T599R mutant cardiomyocytes indeed show increased maximal respiration, spare respiratory capacity, and oxidative metabolic potential. However, they also showed increased non-mitochondrial oxygen consumption and proton leak (Figure 3C-F, Supplementary Figure 5). Overall, the BRAF mutation resulted in an increase in mitochondrial pools and capacity, providing no clarity regarding the inability of these cardiomyocytes to meet a higher ATP demand during acute stress.

**Figure 3.**
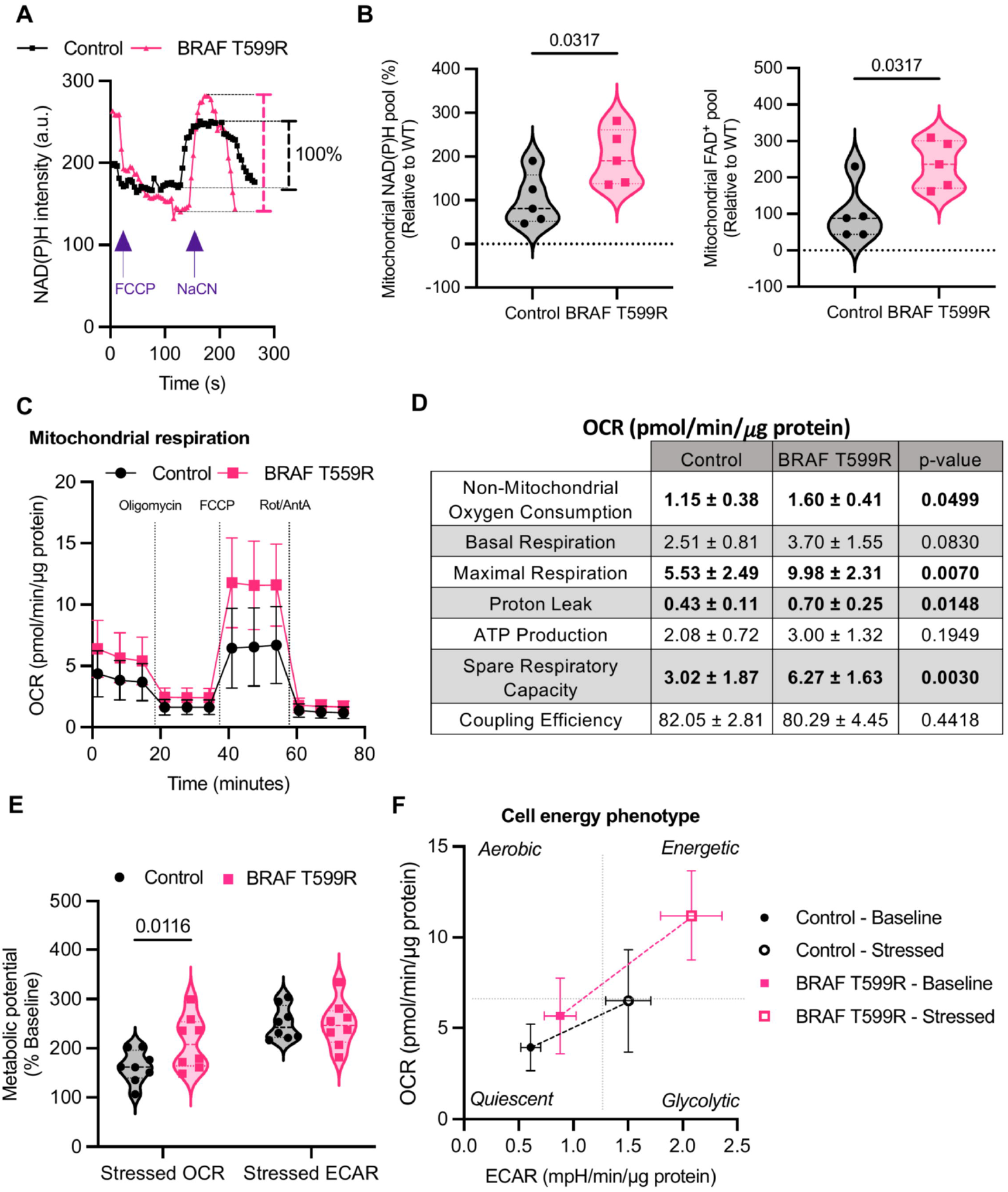
Energetic disturbances in BRAFT599R cadiomyocytes are not due to a decreased mitochondrial capacity. A) Representative NAD(P)H autofluorescence curves from control and BRAF-mutant cardiomyocytes exposed to FCCP (5 **µM)**, and cyanide (4 mM) from the experiment described in Figure 2.F illustrating the calculation of the mitochondrial NAD(P)H pool (B). FAD+ maximally oxidizable mitochondrial pool was calculated similarly. N=5 independent experiments from 2 differentiations. C) Average oxygen consumption rate (OCR) during the mitochondrial respiration stress test measured with the Seahorse XF analyzer. Control and BRAF-mutant CMs were exposed to the ATP synthase inhibitor oligomycin (5 **µM)**, the uncoupling agent FCCP (1 **µM)**, and the respiration complex inhibitors rotenone (1 **µM)** and antimycin A (1 **µM)**. Rates are normalised to total protein measured by Bradford assay. D) Mitochondrial respiration parameters determined from the OCR profile presented as mean ± standard deviation. E) Metabolic potential of control and BRAF-mutant CMs estimated as the percentage increase between stressed (FCCP exposed) and basal OCR and extracellular acidification rate (EGAR). F) Cell energy phenotype based on the normalised OCR and EGAR values in baseline and stressed conditions. Quadrants were set arbitrarily to indicate the direction of bioenergetic changes. N=8 differentiations. Data presented as mean ± standard deviation. For others, violin plots show the distribution, median, and quartiles of the data. The statistical analysis used was a two-tailed Mann-Whitney test except where otherwise specified. p-value < 0.05 considered significant and displayed.

### Mavacamten restores the energy deficiency in BRAF T599R cardiomyocytes

Mavacamten is a myosin inhibitor that was recently approved for the treatment of obstructive hypertrophic cardiomyopathy in adults (36). It has been shown to attenuate hypercontractility by selectively inhibiting the cardiac myosin ATPase at the myofilament (Figure 4A). Therefore, we treated BRAF-mutant cardiomyocytes with mavacamten to evaluate if the heightened energy demand related to myocyte hypercontractility was responsible for the energetic imbalance. Acute exposure to the drug resulted in a small, non-significant reduction in the redox index of the cardiomyocytes determined by NAD(P)H and FAD^+^ autofluorescence, which could indicate a reduction in mitochondrial overactivity (Figure 4B). However, treating the cardiomyocytes during the incremental pacing protocol led to a significant decrease in the ATP fall, indicating, at least, a partial restoration of the energetic balance in the BRAF T599R mutant cardiomyocytes (Figure 4C-D). These results confirm that cardiomyocyte hypercontractility is a major pathological mechanism underlying the metabolic alterations in BRAF-induced RASopathy and that mavacamten improves the energy deficit it produces (Figure 4D-E).

**Figure 4.**
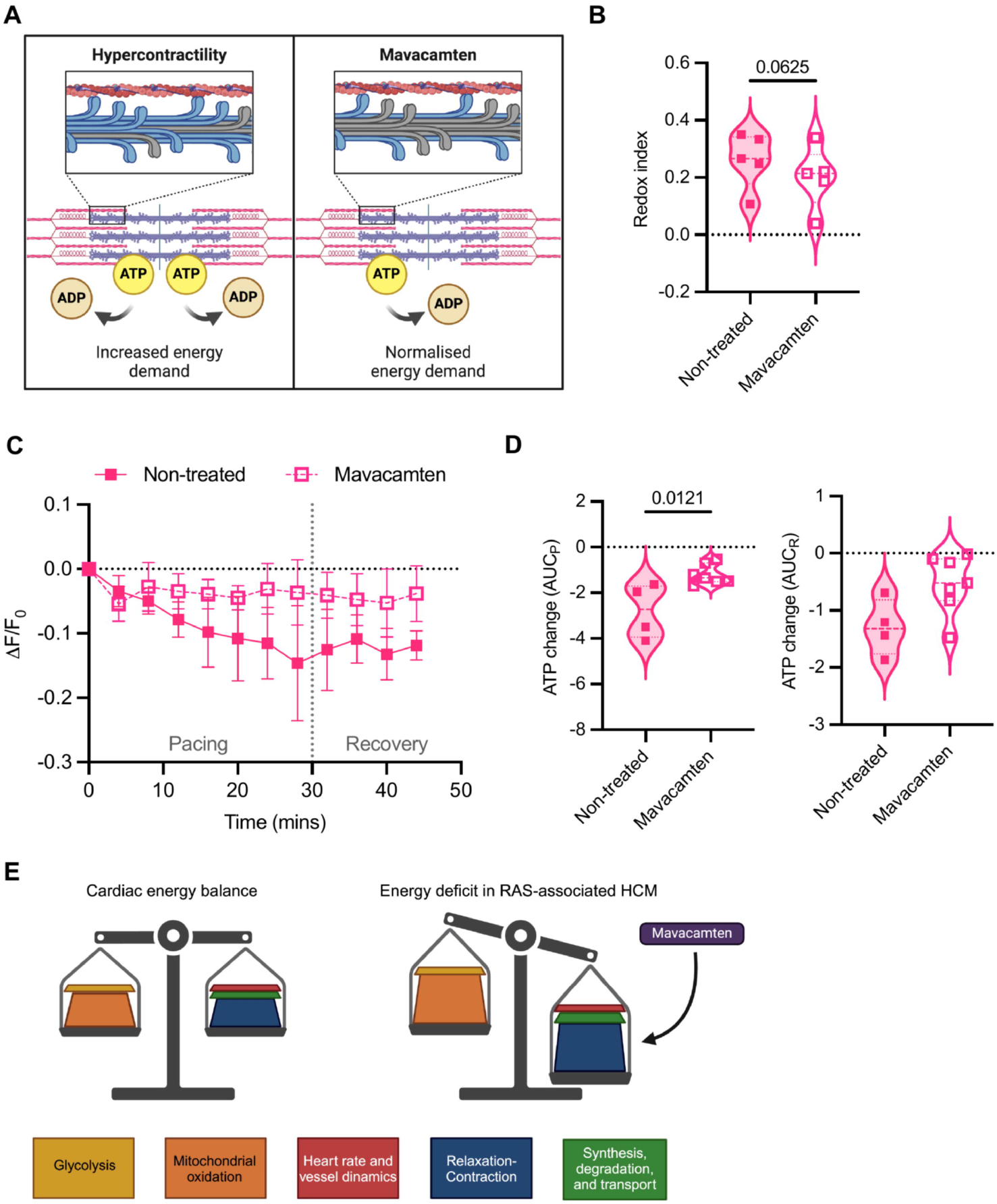
Mavacamten restores the energy deficiency in BRAFT599R cardiomyocytes. A) Effects of mavacamten in sarcomere dynamics and ATP consumption, adapted from Nag *et al*. (20). B) Redox index changes in BRAF mutant CMs calculated from NAD(P)H and FAD+ autofluorescence upon application of mavacamten (0.5 ***µ*M)** as described in Figure 2F-G. N=5 independent experiments from 2 differentiations. Paired Wilcoxon test. C) ATPtracker intensity profile following the electrical pacing protocol described in Figure 2D with mavacamten (0.5 ***µ*M)** treatment. Data presented as mean ± standard deviation. D) Area under de curve (AUG) quantification for the pacing and recovery periods separately. N=4-7 differentiations. Violin plots show the distribution, median, and quartiles of the data. The statistical analysis used was a two-tailed Mann-Whitney test. p-value < 0.05 considered significant and displayed. E) Energy supply and demand balance in the healthy heart and RAS-associated HCM (52).

## Discussion

In recent years the iPSC-CMs have emerged as a powerful platform for disease modelling and therapeutic screening (37, 38). In this study, we characterised an in vitro model of RAS-HCM using *BRAF* p.Thr599Arg mutant cardiomyocytes, a RASopathy-associated mutation. Through a combination of morphological, functional and metabolic analyses we have identified hallmarks of HCM, including hypertrophy, hypercontractility and energetic deficiency under acute stress. Interestingly, treatment with mavacamten ameliorated the energy deficiency, highlighting its potential as a therapeutic agent in RAS-HCM.

### BRAF-mutant cardiomyocytes reflect the pathophysiology of a RASopathy

RAS-HCM shares phenotypic similarities with sarcomeric HCM, including cardiomyocyte hypertrophy, contractile abnormalities, and arrhythmia susceptibility. So far, sarcomeric HCM has been the most widely studied in vitro, recapitulating almost all of the above-mentioned characteristics (39, 40), but hypertrophy is not always detected (41). In this study, we demonstrated that BRAF-mutant cardiomyocytes exhibit significant hypertrophy, which aligns with our and others’ prior studies (11, 12, 26). In addition, the observed upregulation of protein translation pathways and extracellular matrix gene expression follows the transcriptional changes observed in the early stages of cardiac hypertrophy (33), and the increased expression of MYL2 mirrors the expression and correlation of this gene with increased cell size as detected in HCM patient samples (34).

While sarcomeric mutations directly alter the function of the contractile apparatus, resulting in hypercontractility (10, 40), the molecular mechanisms linking RAS-MAPK pathway activation to hypercontractility are indirect and remain incompletely understood. RASopathy mutations, including BRAF (13), MRAS (12), and LZTR1 (42) variants, have been shown to induce calcium-handling defects. It was already reported that BRAF-mutant hiPSC-CMs display increased calcium transient amplitude and a larger pool of stored calcium driving hypercontractility (11). Proposed mechanisms involved include the downregulation of L-type calcium channels and SERCA (43), dysregulation of miRNAs (26), and the crosstalk between the Rho and RAS-MAPK signaling pathways regulating actin-myosin interaction, which has been studied in the setting of cancer and cellular motility (44). It remains to be explored which of these are the main drivers of the changes in RAS-HCM; however, the mechanisms contributing to the hypercontractile phenotype are likely to be multifactorial.

### Energetic vulnerabilities under acute stress

The inability of BRAFT-mutant cardiomyocytes to maintain ATP levels during pacing highlights a critical energetic vulnerability. While basal ATP levels were slightly reduced, the sharp decline during stress reflects an imbalance between energy production and consumption. Interestingly, mitochondrial content and capacity do not appear to be limiting factors. On the contrary, BRAFT-mutant cells exhibited higher maximal respiration rates, and mitochondrial NAD(P)H and FAD+ pools suggest an increased mitochondrial mass. This is not entirely unexpected, as the RAS-MAPK pathway is known to promote mitochondrial respiration, glycolysis, and mitophagy in cancer cells through several pathways, including AKT and AMPK (45, 46). Mitochondrial biogenesis regulation has also been observed in cardiomyocytes through the MAPK-ERK5-PGC1a axis in diabetic cardiomyopathy (47).

It follows, therefore, that the observed energetic deficits could be due to mitochondrial inefficiencies, elevated ATP consumption from hypercontractility, or both. As evidenced by the altered redox ratios during pacing and the increases in non-mitochondrial oxygen consumption and proton leak, it is possible that cumulative mitochondrial damage (48) or oxidative stress (49) could hinder the energetic compensatory response to the ATP demand from the rapid pacing in BRAF-mutant cardiomyocytes. Such inefficiencies may result from chronic endogenous stress linked to the persistent activation of RAS-MAPK signalling. On the other hand, hypercontractility itself constitutes a substantial stressor for cardiomyocytes since ATP consumption follows the same dynamic as contraction (50), meaning that BRAF-mutant cells must match the constantly high ATP demand. Further research is required to determine if mitochondrial damage exists in these cells; however, the recovery observed with the myosin inhibitor suggests the high ATP demand is the major factor contributing to the energy deficiency.

### Mavacamten restores energetic balance

Mavacamten, a small-molecule myosin inhibitor, demonstrated significant efficacy in mitigating the energetic disturbances in BRAF-mutant cardiomyocytes. The mechanism by which mavacamten achieves these effects likely involves modulation of myosin-actin interactions, which are exaggerated in hypercontractile states (20). In sarcomeric HCM, mavacamten rescued abnormal contractility and relaxation by promoting the super-relaxed state of myosin, restoring Ca2+ sensitivity, and ATP consumption (51). These mechanisms appear equally applicable to alleviate the contractile burden in RAS-HCM, broadening the therapeutic range of myosin inhibitors.

While our study provides important insights into the pathophysiology of RASopathy-associated HCM and the therapeutic potential of mavacamten, certain limitations should be acknowledged. The iPSC-CM model does not address different cell types or systemic responses. Furthermore, the precise contributions of oxidative stress, calcium mishandling, and mitochondrial damage remain unclear. Regarding therapeutic development, the experimental design focuses on an acute response. Long-term studies are required to evaluate the durability of its therapeutic effects and potential off-target impacts.

Our findings demonstrate that BRAFT599R cardiomyocytes recapitulate the key features of RASopathy-associated HCM, including hypertrophy, hypercontractility, and energetic dysfunction. Mavacamten effectively restores energetic balance, underscoring its potential as a therapeutic agent for RAS-associated HCM. Future studies integrating patient-specific models and in vivo systems will be essential to evaluate further and refine this therapeutic strategy for a clinical setting.

## Acknowledgements

We would like to express our gratitude to the PARCC’s technical facilities, particularly the iPSC platform. We thank Yunling Xu from the Microscopy platform and Camille Knops from the Flow Cytometry platform for their invaluable support.

## Data availability statement

The data underlying this article are available in the article and in its online supplementary material. The mRNA sequencing data set is available upon request to the corresponding author.

