## Supplementary figures for "Mavacamten improves energy balance in a pre-clinical model of RASopathy-associated hypertrophic cardiomyopathy"

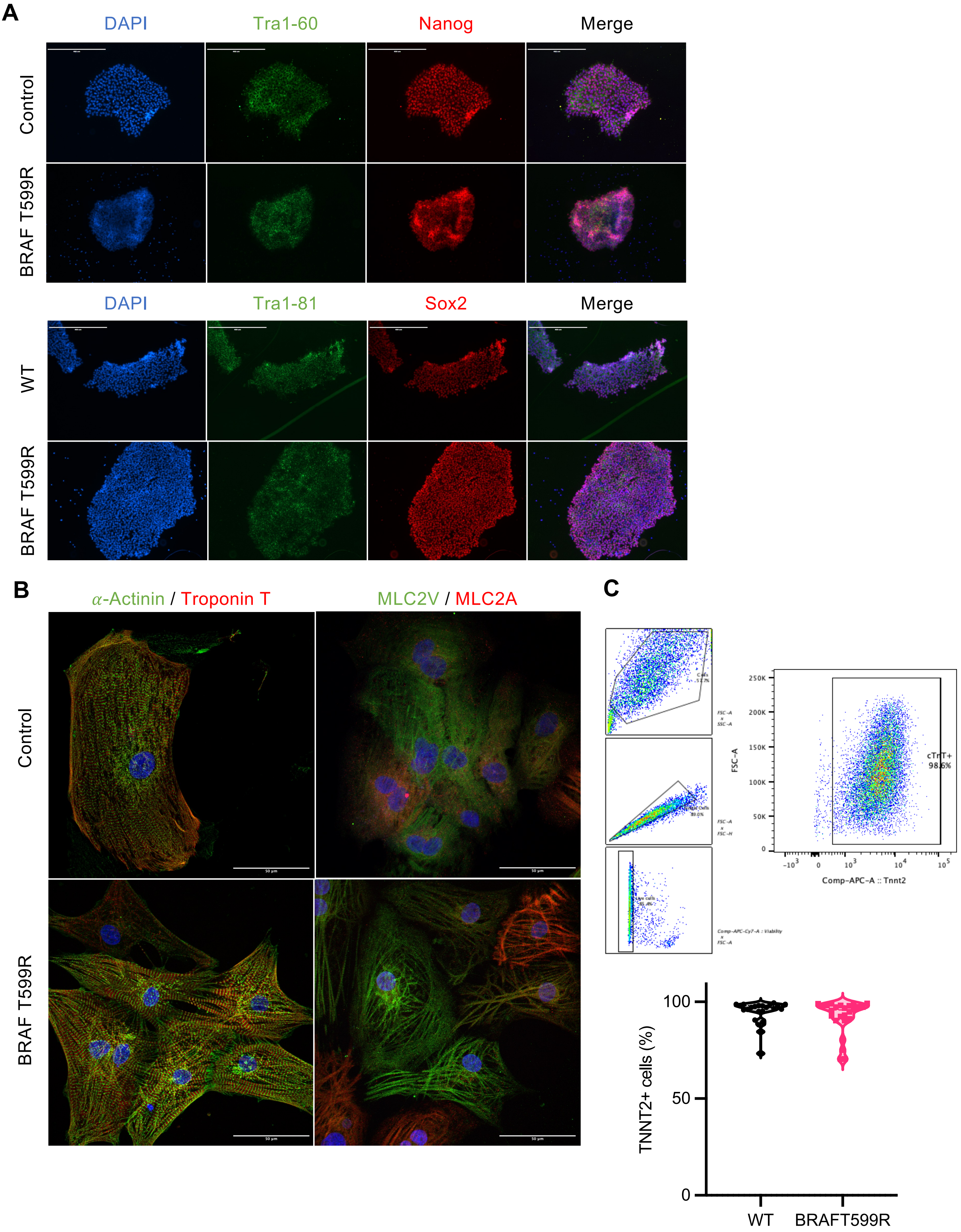


**Supplementary Figure 1. WT and BRAFT599R hiPSC pluripotency markers and hiPSC-CM characterisation.** A) Pluripotency markers detected in WT and BRAF-mutant hiPSC by immunofluorescence (scale bar = 400 $\mu$m). A) Immunofluorescent confocal images of cardiac markers detected in WT and BRAF-mutant CMs (scale bar = 50 $\mu$m). C) Determination of cardiac differentiation efficiency by flow cytometry with the Zombie viability dye and Troponin (TNNT2)-APC coupled antibody. The gating strategy used in the FlowJo analysis software is displayed with the violin plot of the final results. Only differentiations with more than 80% TNNT2+ cells were used for experiments. N=23-25 differentiations.

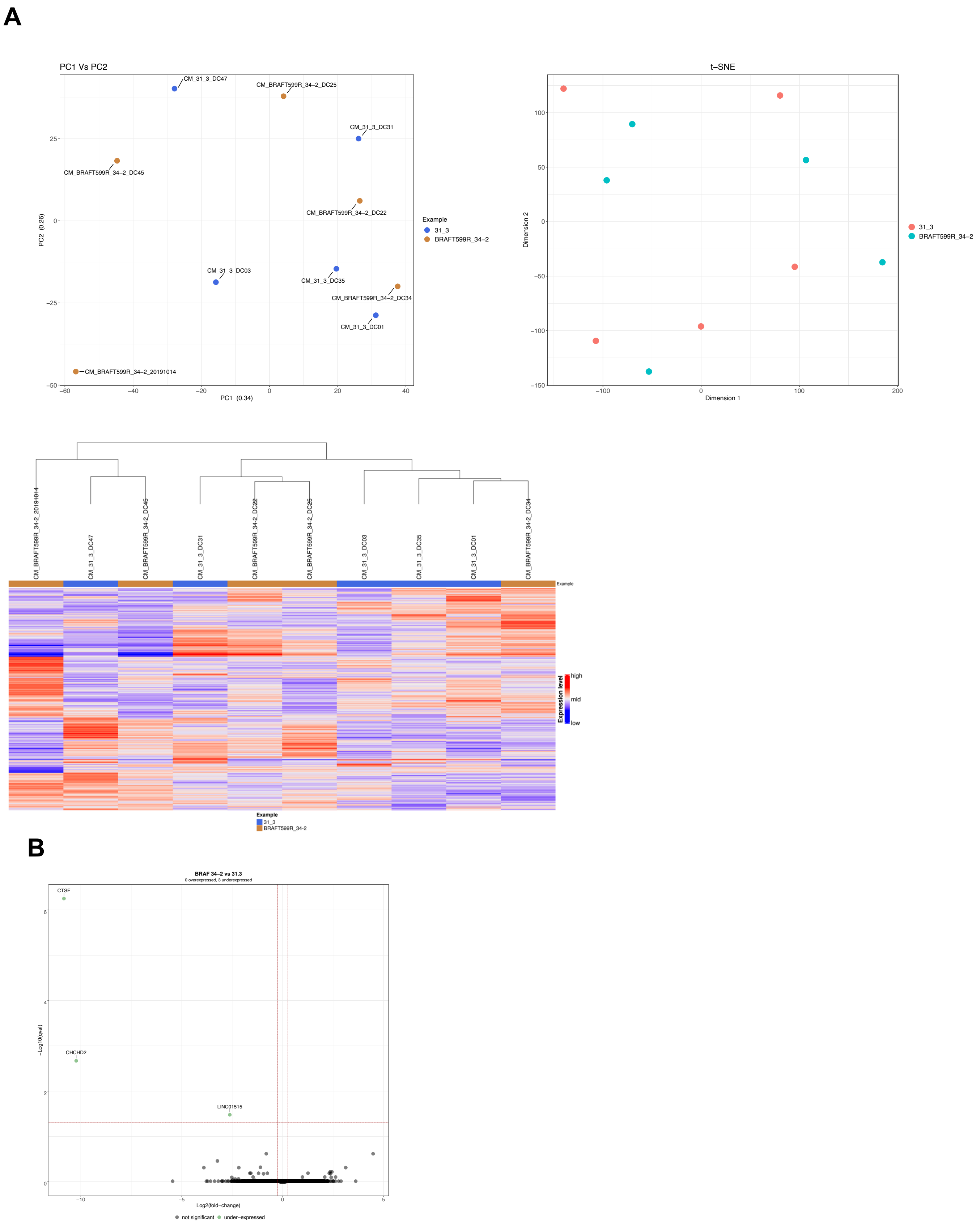


**Supplementary Figure 2. RNA-sequencing analysis.**

A) Principal component analysis (PCA), t-distibuted stochastic neighbor embedding (t-SNE) plots, and hierarchical clustering of the1000 most variable genes from the bulk RNA-sequencing data. The WT CMs are labeled as 31.3. B) Volcano plot of differentially expressed genes in BRAF-mutant CMs vs WT. N=5 differentiations. *CTSF*: Cathepsin F, *CHCHD2*: Coiled-coil-helix-coiled-coil-helix domain containing 2, *LINC01515*: long intergenic non-protein coding RNA 1515.

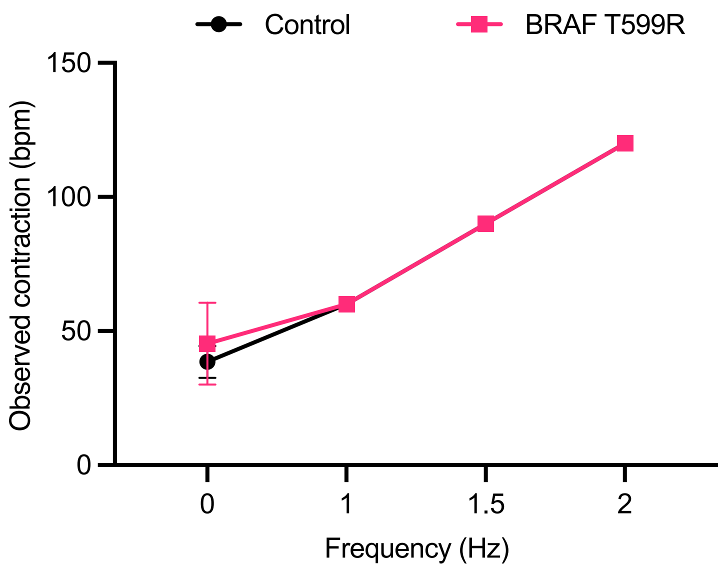


**Supplementary Figure 3. Observed contraction rate with pacing.**
Contraction rate of WT and BRAF-mutant CM observed with increasing pacing frequency. N=3-4 differentiations. Data presented as mean ± standard deviation.

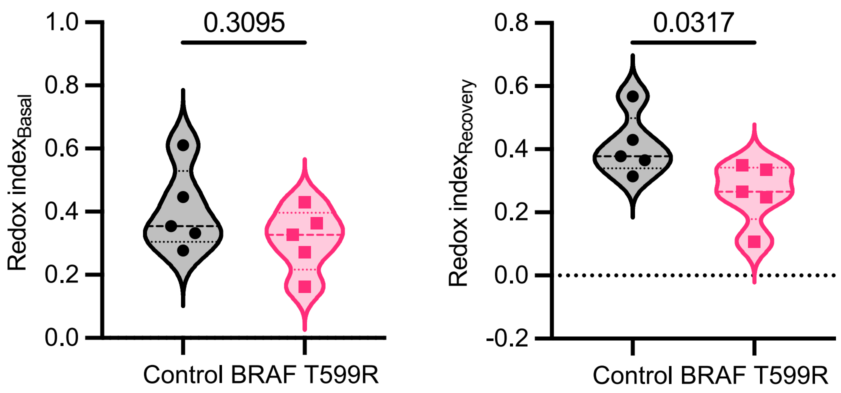


**Supplementary Figure 4. Redox index before and after pacing.**
Redox index in WT and BRAF-mutant CMs in basal condition and after the incremental electrical pacing protocol (Recovery) calculated from NAD(P)H and FAD+ autofluorescence . N=5 indepent experiements from 2 differentiations. Violin plots show the distribution, median, and quartiles of the data. Statistical analysis used was two-tailed Mann-Whitney test. p-value < 0.05 considered significant and displayed.

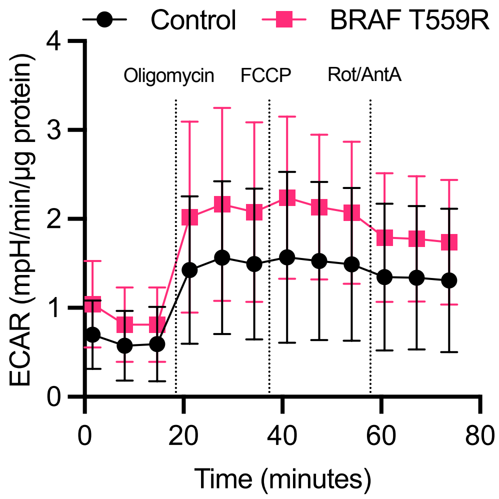


**Supplementary figure 5. ECAR profile from the MitoStress test.**
Extracellular acidification rate (ECAR) profile from the MitoStress test performed to measure mitochondrial respiration. N=8 differentiations. Data presented as mean ± standard deviation.

### MuscleMotion parameters

| | Control | BRAF T599R | p-value |
|---|---|---|---|
| Contraction duration (ms) | **726.1 ± 168.2** | **891.4 ± 114.3** | **0.0176** |
| Time to peak (ms) | 198.4 ± 54.75 | 207.8 ± 63.59 | >0.9999 |
| Relaxation time (ms) | **527.7 ± 146.9** | **683.7 ± 108.9** | **0.0360** |
| 90-to-90 transient (ms) | **726.1 ± 168.2** | **891.4 ± 114.3** | **0.0176** |
| 50-to-50 transient (ms) | **500.6 ± 109.8** | **663.5 ± 119.3** | **0.036** |
| 10-to-10 transient (ms) | 210.2 ± 37.08 | 258.9 ± 90.35 | 0.4735 |
| Contraction amplitude (a.u.) | **287.1 ± 43.11** | **462.7 ± 155.1** | **0.0048** |
| Contraction rate (bpm) | 36.19 ± 11.25 | 32.47 ± 5.35 | 0.4920 |

**Supplementary Table 1. MuscleMotion results.**
Contractility parameters of WT and BRAF-mutant CM monolayers calculated by the MuscleMotion ImageJ plugin and presented as mean ± standard deviation. N=6-9 differentiations. Statistical analysis used was two-tailed Mann-Whitney test. p-value < 0.05 considered significant and highlighted with bold letters.
